# ELIP is preferentially expressed in stomatal guard cells and has a role in stomatal opening upon transition to light

**DOI:** 10.1101/2025.03.26.645183

**Authors:** Dominika Bednarczyk, Maya Cohen, Itzhak Kamara, Vivekanand Tiwari, Eba Abraham Itafa, Preetom Regon, Alon Savidor, Deepanker Yadav, Eduard Belausov, Dana Charuvi

## Abstract

Plant early light-induced proteins (ELIPs) are chlorophyll-binding thylakoid membrane proteins commonly believed to function in photoprotection. However, results of studies with mutants or transgenic plants have been contradictory regarding this function, and the mechanistic workings of ELIPs have mostly remained elusive. We studied: (1) ELIP function in tomato (*Solanum lycoperiscum*) by generation and photosynthetic/physiological characterization of *slelip* mutants under various light conditions; and (2) ELIP tissue/cell localization by monitoring expression of the Venus yellow fluorescent protein fused to the native *SlELIP1* promoter (‘*pSlELIP1*::Venus’) in transgenic tomato. Notably, *slelip* mutants did not display compromised photosynthetic performance and were not more sensitive to photoinhibition compared to parental plants, even following direct sunlight exposure. Surprisingly, *pSlELIP1*::Venus fusions revealed preferential expression in the leaf epidermis and specifically in stomatal guard cells, with no apparent expression in the mesophyll. No major alterations were observed in leaf gas exchange in *slelip* mutants at different light conditions. Intriguingly, stomatal conductance was reduced in *slelip* mutants upon the transition from dark to light. We propose an alternative hypothesis for plant ELIPs as players in the response of guard cells to light, connecting chlorophyll to stomatal function, and presenting a novel research direction for these enigmatic proteins.

## Introduction

Early light-induced proteins (ELIPs) are chlorophyll- and carotenoid-binding proteins that resemble the photosynthetic light-harvesting complex (LHC) antenna proteins. The broader ‘LHC-like’ family encompasses several groups of light-induced thylakoid membrane proteins that do not function in light harvesting: the ‘one-helix protein’ (OHP), the two-helix ‘stress-enhanced protein’ (SEP, also called LIL), the three-helix ELIP and the four-helix PsbS (Adamska, 1997; Montané & Kloppstech, 2000; Yurina *et al*., 2013; Rochaix & Bassi, 2019; Levin & Schuster, 2023; Iwai *et al*., 2024).

Plant ELIPs were discovered as transiently expressed proteins during greening of etiolated *Pisum sativum* (pea) and *Hordeum vulgare* (barley) seedlings (Meyer & Kloppstech, 1984; Grimm & Kloppstech, 1987; Grimm *et al*., 1989), when the developing photosynthetic apparatus is susceptible to photooxidation. Over the years, ELIPs have been shown to be induced (transcripts and in some cases the proteins) in plants at different light and abiotic stress conditions, including red-, blue- and UV light, high light (HL), cold (± HL), heat, salinity, drought and desiccation of resurrection plants (Adamska *et al*., 1992a,b; Adamska & Kloppstech, 1994; Montané *et al*., 1997; Alamillo & Bartels, 2001; Harari-Steinberg *et al*., 2001; Zeng *et al*., 2002; Sävenstrand *et al*., 2004; Peng *et al*., 2008; Floris *et al*., 2013; Oung *et al*., 2022; Wu *et al*., 2024).

Induction of ELIPs has been correlated to the extent of photoinhibition – the accumulation of damage in photosystem II (PSII), and they have been suggested to associate with the PSII reaction center protein D1 and with LHCII fractions at HL (Adamska & Kloppstech, 1991; Adamska, 1997; Heddad *et al*., 2006). ELIPs are also expressed during senescence (Humbeck *et al*., 1994; Binyamin *et al*., 2001; Bhalerao *et al*., 2003; Norén *et al*., 2003) and ripening of tomato fruit (Bruno & Wetzel, 2004), which along with de-etiolation, are plastid developmental processes that involve substantial pigment and metabolic alterations. ELIPs’ expression at all of the above highlights their importance at times when the photosynthetic apparatus is sensitive, and has supported the view that they function in photoprotection (Adamska, 2001; Zeng *et al*., 2002; Yurina *et al*., 2013; Levin & Schuster, 2023). In evergreen trees, (multiple) ELIPs expressed during winter have been proposed to act as quenchers or regulators of sustained thermal dissipation (Zarter *et al*., 2006; Cronn *et al*., 2017; Ye *et al*., 2025).

Results from studies with mutants and transgenic overexpressing plants have been contradictory about the role of ELIPs in photoprotection. Based on the increased photooxidative damage in *Arabidopsis thaliana* plants with reduced ELIP accumulation [in the *chaos* mutant - deficient in the chloroplast signal recognition particle system that targets LHC and ELIP to the thylakoids], and restoration of phototolerance by ELIP overexpression, it was proposed that ELIPs photoprotective role may involve binding of free chlorophyll (Chl) released from turnover of Chl-binding proteins (Hutin *et al*., 2003). Examination of the *A. thaliana* single mutants, *elip1* or *elip2*, showed that their greening tissues had lower Chl content compared to wild type, yet their sensitivity to short-term photoinhibition inflicted by HL or HL and cold was the same as wild type (Casazza *et al*., 2005). The double mutant *elip1/elip2* also showed reduced Chl content in mature leaves and greening seedlings, as well as lower accumulation of the xanthophyll carotenoid zeaxanthin at HL, suggesting that ELIPs may affect the stability or synthesis of photosynthetic pigments (Rossini *et al*., 2006). Nonetheless, *elip1/elip2* plants did not exhibit any significant alterations in their sensitivity to photoinhibition, in non-photochemical quenching (NPQ), in photooxidation, or in the ability to recover from HL stress (Rossini *et al*., 2006). Later on, it was shown that the *elip* mutants are affected in seed germination (Rizza *et al*., 2011). Studies with overexpression of ELIPs have proposed they are involved in regulation of Chl biosynthesis (Tzvetkova-Chevolleau *et al*., 2007; Wang *et al*., 2022). ELIPs have also been connected to Chl biosynthesis during greening via their regulation by the carotenoid-promoting ORANGE protein and the TCP14 transcription factor (Sun *et al*., 2019). Heterologous overexpression of *ELIPs* demonstrated increased tolerance to abiotic stresses (Zhuo *et al*., 2013; Xiao *et al*., 2023), supporting a photoprotective function. Considering the distinct results from different studies, species and conditions, there is still uncertainty about the role(s) of plant ELIPs.

In the present work, we studied ELIPs in tomato (*Solanum lycoperiscum*) plants along two directions. First, by generation of *slelip* mutants and studies of their photosynthetic performance at different light conditions. Second, by investigation of SlELIP1 localization in transgenic tomato plants expressing the Venus yellow fluorescent protein fused to the native *SlELIP1* promoter. Studies with *slelip* mutants indicated that SlELIPs are not required for photoprotection under HL stress. Surprisingly, *pSlELIP1* conferred preferential expression in the leaf epidermis, most predominantly in stomatal guard cells. In line with these results, we found that *slelip* mutants exhibited reduced stomatal conductance upon the transition from dark to light.

## Materials and Methods

### Plants, growth and light experiments

Tomato (*Solanum lycoperiscum* cv. M82) seedlings were grown in soil at long-day conditions (16 h light/8 h dark) at 24/21°C (day/night) under 60-100 µmol photons m^-2^ s^-1^, referred to as low light (‘LL’). For high light (‘HL’) treatment, 7-day old seedlings or 6-week old plants were transferred to a growth chamber under 1000-1200 µmol photons m^-2^ s^-1^. Greenhouse (GH)-grown plants were subjected to sunlight (1500-1800 µmol photons m^-2^ s^-1^). Subsequent recovery (‘REC’) was carried out at initial LL or GH conditions. For greening experiments, seeds were germinated on solid Nitsch medium (Duchefa Biochemie, The Netherlands) without sucrose and grown in the dark for 6 days. On the 7^th^ day, etiolated seedlings were exposed to continuous light (∼150 µmol photons m^-2^ s^-1^). For fruit and seed collection, plants were grown in a GH, with light intensities ranging between 50 to 450 µmol photons m^-2^ s^-1^ along the day. For analysis of the shoot apex, seeds were germinated in a controlled chamber with a light/dark regime of 10/14 h under ∼250 µmol photons m^-2^ s^-1^ or ∼750 µmol photons m^-2^ s^-1^ for 10-12 days. Photon flux densities were recorded using an LI-250A quantum sensor (LI-COR, USA).

### Molecular biology and sequences

Genomic DNA was isolated using the GenElute Plant Genomic DNA Miniprep Kit (Sigma-Aldrich). Complete genomic sequences of ‘*SlELIP1’* (Solyc09g082690) and ‘*SlELIP2’* (Solyc09g082700) were amplified using specific primers. PCR was performed using the DreamTaq Green PCR Master Mix (ThermoFisher). Sequencing was carried out by Macrogen, Inc. Total RNA was extracted using the Trizol reagent (BioLabs, Israel). First-strand cDNA was synthesized using the Verso cDNA Synthesis Kit (ThermoFisher). The reaction included RT Enhancer to prevent genomic DNA carryover, eliminating the need for a separate DNase I treatment. qRT-PCR analysis was performed with the StepOne Plus system (Applied Biosystems) using SYBR Green detection (Applied Biosystems). The *SlSAND* gene [Solyc03g115810; (González-Aguilera *et al*., 2016)] and/or *SlCAC* gene [Solyc08g006960, (González-Aguilera *et al*., 2016)] were used as reference genes, as noted in the figure legends. The list of all primers used is provided in Table S1.

The cleavage sites of the SlELIPs chloroplast transit peptides were predcited by ChloroP (Emanuelsson *et al*., 1999). Transmembrane regions were predicted by TOPCONS (Bernsel *et al*., 2009). Sequence alignments were generated with MultAlin (Corpet, 1988) or Clustal Omega (Madeira *et al*., 2024) and visualized using GeneDoc (Nicholas & Nicholas, 1997).

### Generation of mutants and transgenic plants

#### Vectors

Single guide RNAs (sgRNAs) targeting exons 1, 2 and 3 of *SlELIP1*, and exons 1 and 2 of *SlELIP2* (Table S1) were designed and checked for off-targets using Ensembl Plants (http://plants.ensembl.org). sgRNAs were cloned into a construct containing the *Streptococcus pyogenes* Cas9 (SpCas9) gene using the Golden Braid (GB) system (Sarrion-Perdigones *et al*., 2014). Cas9 was expressed under the constitutive Ubiquitin4-2 promoter from *Petroselinum crispum* (PcUbi4-2) and the Pea3A terminator (Fauser *et al*., 2014). The sgRNAs were expressed under the control of *A. thaliana* U6-26 RNA Pol III promoter and the Kanamycin resistance gene (Kana) was expressed under the Nopaline Synthase (Nos) promoter and Nos terminator.

The *SlELIP1* promoter (‘*pSlELIP1’*) sequence was based on NCBI GenBank accession no. MK867692.1 (Timerbaev & Dolgov, 2019). The corresponding sequence (2165-bp upstream of the translation start site) was retrieved from the Solanaceae Genomics Network (https://solgenomics.net/; reference genome SL4.0). Alignment to MK867692.1 is shown in File S1. *pSlELIP1* was synthesized by GenScript Inc. (USA) and fused to the coding sequence of Venus yellow fluorescent protein (‘*pSlELIP1*::Venus’) or to the *SlELIP1* coding sequence followed by Venus (‘*pSlELIP1*::SlELIP1-Venus’).

#### Plant transformation and genotyping

Plasmids were transformed into *Agrobacterium tumefaciens* (strain GV3101) by electroporation, and selected on LB plates with 200 μg mL^-1^ spectinomycin and 50 μg mL^-1^ gentamycin. T0 cotyledons of tomato (cv. M82) were transformed with the transformed *Agrobacterium*, and then passed through several selection phases according to (Dahan-Meir *et al*., 2018). All media and hormones were from Duchefa Biochemie (The Netherlands), aside from Indole-3-butyric acid (Sigma-Aldrich). Two to three small tomato leaflets were collected from plantlets of the T0 generation, frozen in liquid nitrogen, and kept in −80℃ for further analysis. Verification of transgenic T0 plants (Kana resistant) was done by PCR amplification of the Cas9 sequence from genomic DNA. Cas9-positive transformants were transferred to soil and grown in greenhouse conditions to obtain T1 seeds from self-pollinated T0 plants. T2 seeds were obtained by an additional round of seed collection from T1 self-pollinated plants. For genotyping, genomic DNA was isolated from T1 and T2 plants and the regions spanning the complete genomic region of *SlELIP1* and *SlELIP2* were amplified in order to identify the targeted sites. Examination of T2 homozygous plants showed that all of them passed the same *SlELIP1/2* mutations from T1 to T2 generation, without any further mutation or revision. Homozygous T3 plants were used for the experiments.

### Chlorophyll-*a* fluorescence

Pulse-amplitude-modulated (PAM) Chl-*a* fluorescence recordings were conducted on detached leaves (from 6-week-old plants exposed to different light conditions) using a Maxi-PAM imaging system (Heinz Walz GmbH, Germany). Prior to the measurements, plants were dark-adapted for 30 min. Minimal fluorescence (F0) was determined with the measuring light (9 µmol photons m^-2^ s^-1^), and the maximal fluorescence (Fm) was determined by application of a saturating pulse (SP, 6000 µmol photons m^-2^ s^-1^). Following light induction (using 531 µmol photons m^-2^ s^-1^ provided by the MAXI-PAM), light curves were recorded with light steps between 1-926 µmol photons m^-2^ s^-1^. Calculations of the parameters shown: the maximum quantum yield of PSII, Fv/Fm = (Fm-F0)/Fm); PSII operating efficiency, Y(II) = (Fm’-F)/Fm’; Non-photochemical quenching (NPQ) = (Fm-Fm’)/Fm’ (Baker, 2008).

Fv/Fm recordings of plants in the greenhouse or under sunlight were conducted using a portable PAM-2000 fluorometer (Heinz Walz GmbH, Germany). Leaves (attached to the plants) were dark-adapted for 20-30 min using dark leaf clips (DLC-8, Heinz Walz GmbH, Germany), and the ‘Da-2000’ program was used to determine F0 and Fm, and Fv/Fm calculated as above.

### Gas-exchange measurements

Gas exchange recordings of greenhouse-grown plants were conducted using a portable LCi photosynthesis system (ADC BioScientific Ltd., UK) with a clear-top chamber clamping a leaf area of 6.25 cm^2^, at ambient CO_2_ and under different light conditions (greenhouse, sunlight, artificial light), as specified in the figures. For morning measurements, plants grown in a greenhouse were brought into a dark room prior to dawn, and gas exchange recordings were conducted along with initial illumination of the leaves using white LED arrays. All measurements were carried out on attached leaves.

### Liquid chromatography mass spectrometry

For discovery analysis of orange fruit (M82), liquid nitrogen-frozen ground powder of the pericarp was pooled from 5 biological replicates (from different plants). For targeted analysis, frozen powder from orange fruit pericarp was pooled from 3 biological replicates of each genotype (M82, AC21, AC81 and IK6). For discovery analysis of the leaf epidermis vs whole leaf, M82 plants grown at LL were subjected to HL (700-1000 photons m^-2^ s^-1^ for 4 h), followed by immediate collection of epidermal peels and whole leaf samples, freezing in liquid nitrogen and grinding into fine powder. Sample preparation, liquid chromatography and mass spectrometry of fruit samples were carried out as described (Itafa *et al*., 2025), with some modification for leaf samples. All methods are fully detailed in the Supporting Information.

For discovery analysis, raw data was processed with the MetaMorpheus v0.0.313 software (Solntsev *et al*., 2018) for fruit samples, or with Spectronaut v19.2 (Biognosys) for leaf samples. Data was searched against the *S. lycopersicum* protein database (SlELIP1 and SlELIP2 sequences included) as downloaded from Uniprot (www.uniprot.com) appended with common protein contaminants. Enzyme specificity was set to trypsin and up to two missed cleavages were allowed. Carbamidomethylation of cysteines was set as a fixed modification, and oxidation of methionines was set as a variable one. Protein identifications were filtered at q-value < 0.01. The minimal peptide length was 7 amino-acids. Peptide identifications were propagated across samples using the match-between-runs option checked. For targeted analysis, raw data were loaded into the Skyline software (MacLean *et al*., 2010), peak boundaries and relevant transitions were manually refined, and peak intensities were calculated.

### Confocal microscopy imaging

Freshly cut pieces or cross-sections of cotyledons or mature leaves were imaged with a Leica SP8 laser scanning microscope system (Leica, Wetzlar, Germany), using an HC PL APO CS2 20x/0.75 objective or an HC PL APO CS 63x/1.2 water immersion objective (Leica, Wetzlar, Germany), and the Leica Application Suite X software (LASX). Fluorescence images were acquired following excitation with a solid state laser (λ_ex_ = 514 nm) operating at 2.3% intensity. Emission of Venus Yellow Fluorescent Protein was collected using an HyD detector at the range of 520-560 nm, and emission of chlorophyll was collected using a PMT detector at the range 660-760 nm. *Z*-sections were acquired at 0.6847- or 0.3557-µm steps for 20x or 60x, respectively. Image processing for figure preparation was done using the Fiji open-source package (Schindelin *et al*., 2012). Brightness and contrast adjustments, for improved visualization, were applied to whole images uniformly.

### Statistics

Statistical analysis was conducted with the software JMP Version 16 (SAS Institute Inc., NC, USA). Differences between mutant plants and M82 plants were assessed using the Student’s T-test, and are labelled using asterisks, with the significance level detailed in the figure legends.

## Results

### Expression of the early light-induced proteins (ELIPs) in tomato

Tomato (*S. lycoperiscum*) plants possess two *ELIP* genes (Fig. 1), whose coding sequences are found in tandem in the genome: Solyc09g082690, which we term ‘*SlELIP1*’, and Solyc09g082700, termed ‘*SlELIP2*’ (Fig. 1a). *SlELIP2* lacks the second exon, and accordingly a stretch of 24 amino acids from the polypeptide sequence, in proximity to the chloroplast transit peptide cleavage site (Fig. 1b). SlELIPs possess conserved chlorophyll-binding LHC motifs in the first and third transmembrane helices (Fig. 1b, and see alignment to the *A. thaliana* ELIPs in Fig. S1).

**Fig. 1.**
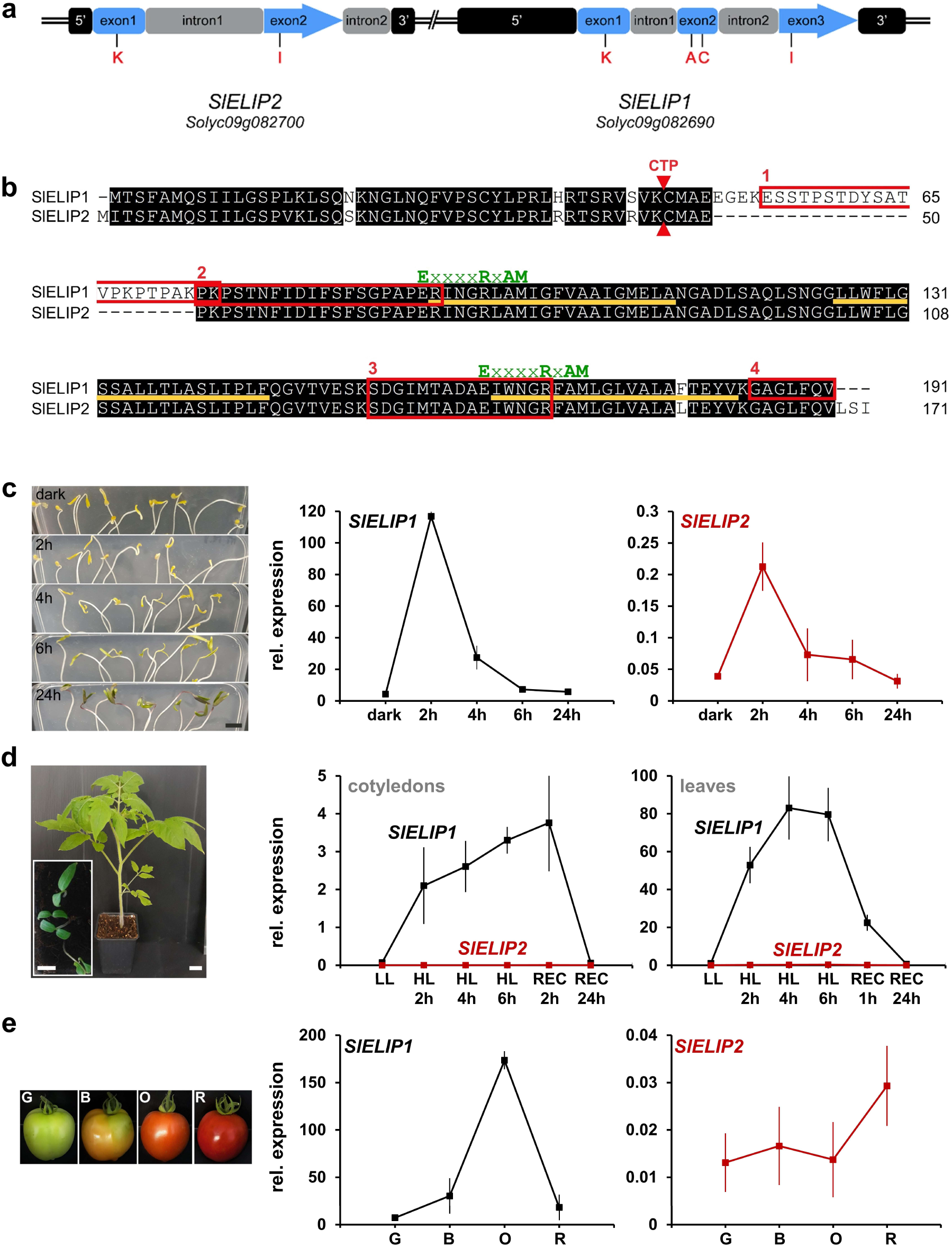
ELIPs in *S. lycopersicum* and their expression at different developmental stages. **(a)** Schematic representation of the *SlELIP1* (Solyc09g082690) and *SlELIP2* (Solyc09g082700) genomic region; ‘**//**’ represents ∼3 kb; Red letters denote gRNAs used for targeting the genes. **(b)** Alignment of the SlELIP1 and SlELIP2 polypeptides. Red triangles mark the predicted cleavage site of the chloroplast transit peptide (CTP). Orange lines indicate the predicted transmembrane regions. The chlorophyll-binding LHC motifs, ‘ExxxxRxAM’, are noted in green. Red numbered boxes show the SlELIP peptides identified by LC-MS/MS of orange fruit. **(c-e)** Expression of *SlELIP1* and *SlELIP2* (red) assayed by qRT-PCR: **(c)** 7-day-old etiolated cotyledons (‘dark’) exposed to continuous light (∼150 µmol photons m^-2^ s^-1^) for 2-, 4-, 6- and 24 h. Scale bar: 1 cm. **(d)** 7-day-old cotyledons (inset) and leaves from 6-week-old plants [Scale bars: 2 cm and 1 cm (inset)] grown at low light (‘LL’, ∼60 µmol photons m^-2^ s^-1^), then exposed to high light (‘HL’, ∼1000 µmol photons m^-2^ s^-1^) for 2-, 4-, and 6 h, followed by recovery (‘REC’; 1- or 2- and 24 h) under LL. **(e)** Expression in the pericarp of green (G), breaker (B), orange (O) and red (R) fruit. For all experiments, expression was normalized to the reference gene *SlSAND*. Values shown are means ± SD of three biological (plant) replicates, each analyzed with three technical replicates.

The expression of the *SlELIP* genes in different tissues and at different conditions was characterized by qRT-PCR (Fig. 1c-e). First, expression was assayed during the etioplast-to-chloroplast transition (greening) in 7-day-old etiolated seedlings before and after their exposure to continuous light (Fig. 1c). Strong induction (∼30-fold) of *SlELIP1* expression was evident after 2 h of light exposure, followed by a sharp decline after 4 h, and a further decline after 6- and 24 h back to the ‘dark’ level. *SlELIP2* showed a similar pattern, but its expression level was considerably lower than *SlELIP1*. We also found that *SlELIP1* exhibited higher expression compared to *SlELIP2* in the shoot apex (Fig. S2), where the proplastid-to-chloroplast transition commences (Charuvi *et al*., 2012; Dalal *et al*., 2018). The expression of *SlELIP*s in response to changes in light intensity was assessed in 7-day-old cotyledons and leaves from 6-week-old plants grown at low light (LL; ∼60 µmol photons m^-2^ s^-1^), subjected to high light (HL; ∼1000 µmol photons m^-2^ s^-1^), and following recovery (REC) periods of 1- or 2 h and 24 h (Fig. 1d). In cotyledons, *SlELIP1* expression was induced by 30-47-fold during HL exposure (2-, 4- and 6 h). Its expression also slightly increased after 1 h REC, but declined back to the initial LL level after 24 h REC. In leaves, the expression of *SlELIP1* was induced up to 80-fold during HL exposure and then declined after 2 h REC and back to the initial level after 24 h. *SlELIP2* was also induced by HL, but its relative expression was considerably lower (Fig. 1d). *SlELIPs* expression in fruit pericarp was determined at different developmental stages representing the chloroplast-to-chromoplast transition: green (G), breaker (B), orange (O) and red (R) (Fig. 1e). *SlELIP1* was considerably induced during ripening, with a peak increase in O-stage fruit (24-fold higher vs G fruit), while *SlELIP2* expression was quite low and did not change.

Earlier, we detected an SlELIP peptide in leaves from HL-stressed tomato plants by liquid chromatography tandem mass spectrometry (LC-MS/MS) (Bednarczyk *et al*., 2020). Here, we subjected O-stage fruit, which exhibited the highest relative gene expression (Fig. 1e), to LC-MS/MS discovery analysis. Four distinct tryptic peptides (cleavage C-terminally to lysine and arginine residues) corresponding to SlELIPs were detected in the analysis (Fig. 1b, red boxes): three of them, #1, 2 and 4 are unique to SlELIP1, while peptide #3 is shared by both SlELIP1 and SlELIP2. This confirmed the presence of SlELIP1 on the protein level, but did not provide direct evidence about the presence of SlELIP2, whose gene expression was considerably lower in all tissues.

### *slelip* mutants

To study the roles of SlELIPs, specifically SlELIP1, we generated several mutants by CRISPR/Cas9. Three mutants were selected for the current study: two mutants in *SlELIP1* and a double mutant in both *SlELIPs*. The lines are named according to the gRNA(s) used (red letters in Fig. 1a): ‘AC21’ and ‘AC81’ for *slelip1* mutants and ‘IK6’ for the double mutant. The genomic deletions and putative polypeptide sequences aligned to the M82 sequences are respectively shown in Figs. S3 and S4. AC21 possesses deletions of 3 nucleotides (nt) in *SlELIP1* resulting in mutation of several amino acids (a.a.) and one a.a. deletion. AC81 has a deletion of 26 nt in *SlELIP1*, predicted to result in mutation of 3 a.a. and a premature stop codon at position 64. In the IK6 double mutant, *SlELIP1* exhibited a 2-nt deletion and *SlELIP2* a 1-nt deletion; these are predicted to result in mutation of a few a.a. and premature stop codons at the first transmembrane region of both SlElips (Figs. S3 and S4). qRT-PCR of *SlELIP1* and *SlELIP2* in leaves at LL, HL and REC light conditions showed a general reduction in expression in the mutant lines compared to M82 (Fig. S5). Targeted LC-MS/MS analysis of O-fruit pericarp tissue showed that the common SlELIP peptide (#3 in Fig. 1b) was detected in M82, but not in the three mutant lines (Fig. S6).

*ELIP* genes are induced during de-etiolation [e.g. (Meyer & Kloppstech, 1984; Casazza *et al*., 2005), Fig. 1c], and the accumulation of chlorophyll (Chl) during this process has been shown to be delayed in *A. thaliana elip* mutants (Casazza *et al*., 2005; Rossini *et al*., 2006). To assess SlELIP functionality during this process in the tomato mutants, we quantified Chl content in 7-day-old etiolated seedlings of M82 and the AC21, AC81 and IK6 mutant lines at 6- and 24 h after the onset of illumination. Compared to M82, the level of Chl *a*, Chl *b* and Chl *a* + *b* were significantly lower in all three mutants at both time points (Fig. 2a,b), indicating that SlELIPs were non-functional in the mutants.

**Fig. 2.**
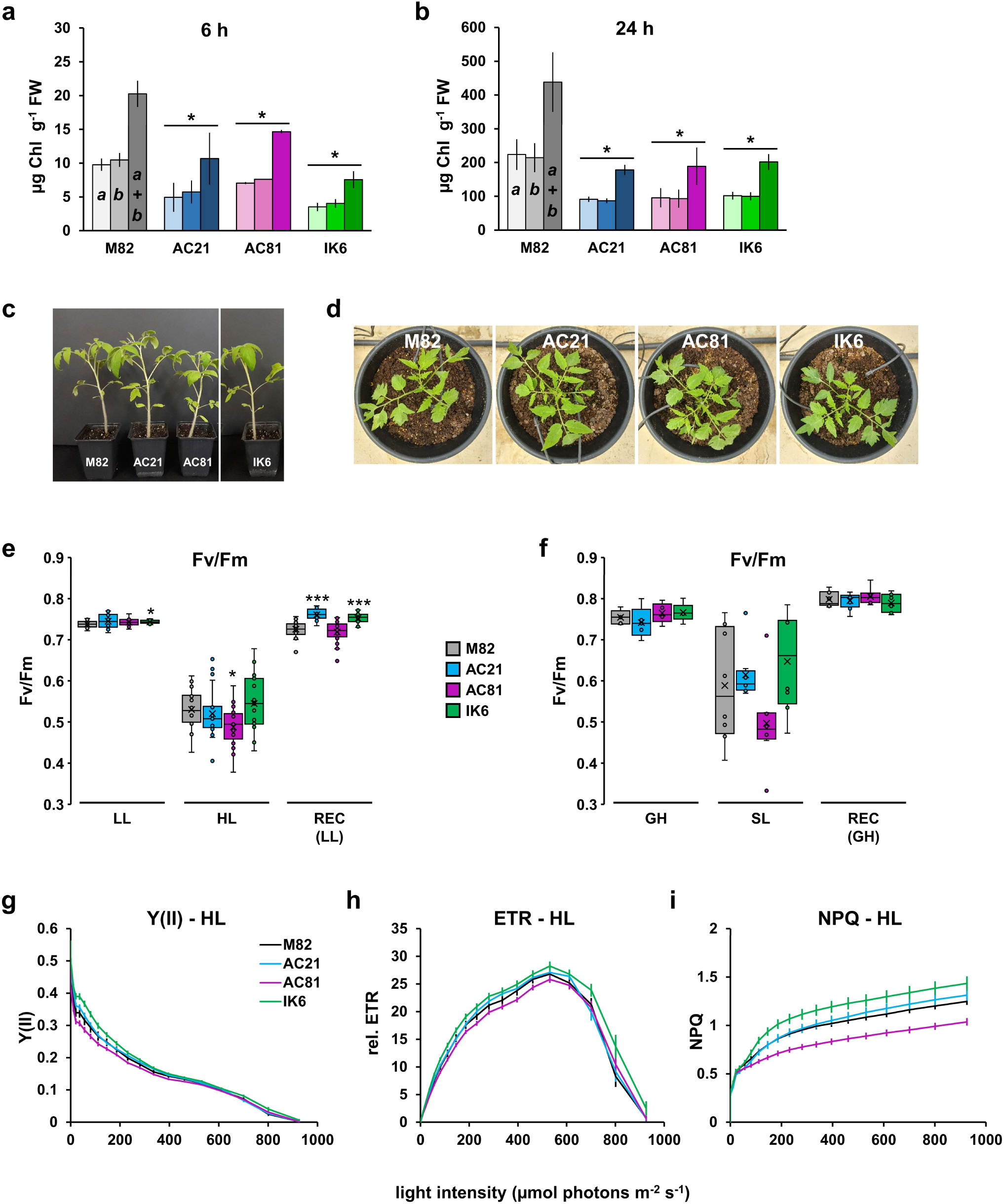
*sleip* mutants exhibit delayed greening, but do not have compromised photosynthetic performance. **(a, b)** Chlorophyll accumulation during greening. Bars depict the levels of chlorophyll (Chl) *a*, Chl *b* and Chl *a* + *b* in 7-day-old etiolated seedlings of M82 and mutant lines AC21, AC81 and IK6 exposed to **(a)** 6- and **(b)** 24 h of continuous white light of ∼150 µmol photons m^-2^ s^-1^. Values are means ± SD of three boxes of 50 seedlings each. Asterisks denote statistically significant differences compared to M82 (*p* < 0.05). Representative pictures of ∼5-week-old plants grown **(c)** in a growth room at low light (LL; ∼60 µmol photons m^-2^ s^-1^), or **(d)** in a greenhouse. **(e-i)** Photosynthetic parameters derived from chlorophyll *a* fluorescence recordings. **(e)** Maximum quantum yield of PSII (Fv/Fm) of plants grown at LL, exposed to high light (HL; 6 h of ∼1200 µmol photons m^-2^ s^-1^) and a recovery period (REC; 24 h at LL). **(f)** Fv/Fm of plants grown in a greenhouse (GH), exposed to direct sunlight (SL; 1500-1800 µmol photons m^-2^ s^-1^ for 6 h), and a recovery period (REC; 24 h in the GH). Significant differences vs M82 at a specific condition are marked \**p* < 0.05, \*\*\**p* < 0.001 (Student’s T-test); number of leaves n ≥ 12 from different plants **(e)**, and n = 5-11 leaves from 3 plants of each genotype **(f)**. Light curves showing **(g)** the effective quantum yield of PSII, **(h)** relative electron transport rate (ETR), and **(i)** non-photochemical quenching (NPQ) of plants after HL exposure. Graphs **(g-i)** show means ± SE, n ≥ 12 leaves, each from a different plant.

### Photosynthetic performance is not considerably altered in *slelip* mutants

The phenotype of 5-week-old mutant plants grown at LL (growth room) or in a greenhouse was similar to M82 plants (Fig. 2c,d). To determine whether SlELIPs are required for a photoprotective role and/or affect photosynthetic performance, we carried out Chl-*a* fluorescence measurements of plants under different light conditions.

First, 6-week-old plants grown at LL were subjected to HL followed by REC, as described in Fig. 1d. To assess the level of photoinhibition, the maximum quantum yield of PSII (Fv/Fm) was determined (Fig. 2e). No major differences were observed in the Fv/Fm values of LL-adapted plants, with the exception of a slightly higher value in IK6 vs M82. Following HL stress, Fv/Fm expectedly declined in all genotypes, with only AC81 exhibiting lower values vs M82. After 24 h REC, Fv/Fm recovered in all plants, and was in fact even higher in AC21 and IK6 compared to M82 and AC81 (Fig. 2e). The same plants were also analyzed by a light curve assessment after HL stress (Fig. 2g-i). No differences between M82 and the mutant lines were noted in the effective quantum yield of PSII (Fig. 2g), or in the relative electron transport rate (Fig. 2h). Some differences in the magnitude of non-photochemical quenching (NPQ) were observed between the genotypes, with AC81 exhibiting relatively lower NPQ and IK6 somewhat higher NPQ (Fig. 2i).

To assay plants adapted to higher light, Fv/Fm was assessed for plants that grew at greenhouse (GH) conditions (∼550 µmol photons m^-2^ s^-1^ at mid-day). Fv/Fm under these conditions was also similar between the genotypes (Fig. 2f, ‘GH’). To assess photoinhibition after light stress, plants were exposed to direct sunlight (‘SL’) on a sunny day for 6 h (1500-2000 µmol photons m^-2^ s^-1^). Fv/Fm values were more variable under these conditions (Fig. 2f, ‘SL’). While the values of AC81 plants were somewhat lower, none of the mutants significantly differed from M82. The same plants exposed to SL were re-measured 24 h later in the GH; in all plants Fv/Fm recovered to the same extent (Fig. 2f, ‘REC’).

### SlELIP1 is preferentially expressed in the leaf epidermis, most prominently in stomatal guard cells

To study the tissue localization of SlELIP1, we generated transgenic tomato lines expressing the native *SlELIP1* promoter [File S1; following the ‘full-length’ promoter described by (Timerbaev & Dolgov, 2019)] fused to the ‘Venus’ yellow fluorescent protein – ‘*pSlELIP1*::Venus’, in short ‘V’. [Venus in ‘V’ lines does not carry a specific intracellular localization sequence, and is therefore expected to spread in nuclei and the cytosol.] Surprisingly, we found that Venus expression is highly preferential to the leaf epidermal layers (Fig. 3). Confocal microscope images showing Chl and Venus fluorescence in leaf cross-sections from LL-grown M82 and plants from independent ‘V’ lines (Fig. 3a-d) revealed Venus expression in adaxial pavement cells (asterisks in Fig. 3b-d), and in abaxial stomatal guard cells (‘GC’; arrowheads in Fig. 3b-d). No Venus fluorescence was visible along the mesophyll layers (Fig. 3b-d). Confocal examination of the two leaf surfaces from the epidermis and into the palisade-(upper) or spongy (lower) mesophyll provided a comprehensive view of Venus expression in the epidermis (Fig. 3e-l). Intriguingly, in both adaxial- and abaxial leaf surfaces, Venus fluorescence was most prominent in stomatal GC (Fig. 3f,h,j,l). The specific expression of *pSlELIP1*-driven Venus in adaxial (but not abaxial) pavement cells of LL-adapted leaves is also apparent in the images (Fig. 3f,h,j,l). No presence of Venus fluorescence aside chlorophyll fluorescence of the mesophyll tissues layers was observed (Fig. 3e,g,i,k). The same pattern of Venus expression was also observed in cotyledons of LL-adapted V plants (shown for 5 lines; Fig. S7).

**Fig. 3.**
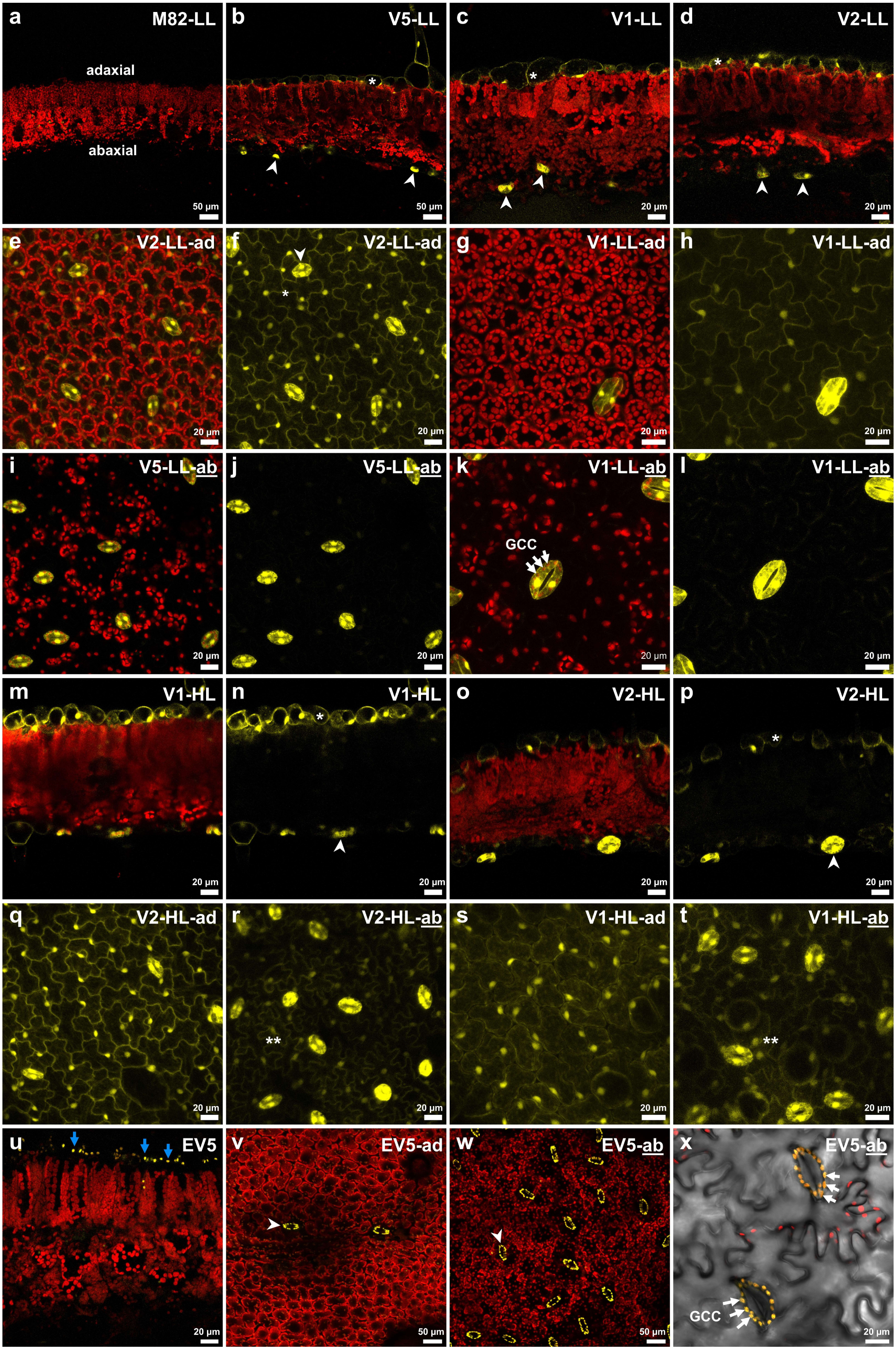
*pSlELIP1* confers expression in the leaf epidermis, particularly in stomatal guard cells. Confocal microscope images showing chlorophyll (Chl; red) and Venus-YFP (yellow) fluorescence in leaf cross-sections and in adaxial (‘ad’) and abaxial (‘<u>ab</u>’) leaf surfaces of plants adapted to low light (‘LL’; 80-100 µmol photons m^-2^ s^-1^), or ones grown at LL and exposed to high light (‘HL’; ∼1000 µmol photons m^-2^ s^-1^). ‘V’ lines (‘V1’, ‘V2’ and ‘V5’) are plant lines expressing *pSlELIP1*::Venus. **(a-d)** Cross-sections showing Chl and Venus in **(a)** M82, and in **(b-e)** V lines. **(e-h)** Views of the adaxial surfaces of V lines showing: **(e, g)** both Chl and Venus, or **(f, h)** Venus. **(i-l)** Views of the abaxial surfaces of V lines showing: **(i, k)** both Chl and Venus, or **(j, l)** Venus. The brightness of images showing only Venus **(f, h, j, l)** was increased to show the comparatively lower fluorescence of pavement cells (where present). **(m-p)** Cross-section images of leaves from V plants exposed to HL, showing: **(m, o)** both Chl and Venus, or **(n, p)** Venus. **(q, s)** Venus fluorescence in adaxial, and **(r, t)** abaxial surfaces at HL. **(u-x)** Chl and Venus fluorescence in ‘EV5’, a line expressing *pSlELIP1*::SlELIP1-Venus: **(u)** leaf cross-section, **(v)** adaxial- and **(w)** abaxial surface. **(x)** Transmission image of the abaxial surface with overlaid Chl and Venus fluorescence. Images of leaf cross-sections are 6-10 µm thick (*Z*-thickness); images of leaf surfaces are 14-37 µm thick (encompassing the epidermal layer and part of the palisade- or spongy mesophyll in adaxial or abaxial surfaces, respectively), aside from panel x (*Z* = 4.3 µm). Arrowheads mark stomata (guard cells); * shows Venus in adaxial pavement cells; ** show Venus in abaxial pavement cells; white arrows point to guard-cell chloroplasts (GCC); blue arrows point to plastids of the adaxial epidermis in EV5.

As the expression of *SlELIP1* is induced by HL (Fig. 1d), V plants were subjected to HL (∼1000 µmol photons m^-2^ s^-1^) in order to determine if the expression pattern changes, and specifically whether it is induced in the mesophyll tissues. In fact, HL did not induce Venus expression in the mesophyll (Fig. 3m-p). Interestingly, HL induced Venus expression in abaxial pavement cells, visible in the leaf surface images (two asterisks in Fig. 3r,t). The distinct induction of expression in abaxial pavement cells, yet apparent lack of induction in mesophyll cells, points to an epidermal-specific response.

To visualize the tissue localization along with the cellular localization of SlELIP1, expected to reside in chloroplasts, we also generated plants expressing *pSlELIP1*::SlELIP1-Venus, termed ‘EV’. Imaging leaf cross-sections of ‘EV’ revealed that Venus fluorescence localizes to small chloroplasts of the adaxial epidermis (blue arrows in Fig. 3u). Furthermore, examination of the adaxial and abaxial leaf surfaces clearly showed Venus expression in GC (arrowheads in Fig. 3v,w), in line with the results from V lines. A detailed view of GC of the abaxial epidermis showed the specific co-localization of Venus with chlorophyll of GC chloroplasts (arrows in Fig. 3x).

To confirm the microscopy results, leaf (upper) epidermal peels and whole leaf samples from M82 plants exposed to HL were analyzed by discovery LC-MS/MS. ∼7500 proteins were identified in the analysis (Table S3), of them we found 28 LHC proteins, also called chlorophyll *a*/*b* binding proteins (CABs), as well as the ‘light-harvesting-like’ proteins ELIP1, OHPs, SEPs/LILs and PsbS (Table S2). For all of these, we examined the ratio between the upper epidermis vs the whole leaf. Notably, SlELIP1, identified with 6 peptides, was found to have a ratio of 6.3, while for the other 27 proteins the epidermis vs leaf ratio ranged between 0.6 and 2.1 (Table S2). We also examined the intensity-based absolute quantification [iBAQ, (Schwanhüusser *et al*., 2011)] values, which showed that SlELIP1 is as abundant as some of the other CABs in the epidermis or leaf (Table S2), and is in fact amongst the top 10% most abundant proteins in the upper epidermis at HL (Table S3). These results confirm the notable enrichment of the native SlELIP1 in the epidermis at HL, and highlight this unique characteristic of SlELIP as compared to other LHC and LHC-like proteins.

### Stomatal conductance is reduced in *slelip* mutants upon the transition from dark to light

Following the finding that SlELIP1 is expressed in the epidermis, and prominently in stomatal GC, we investigated whether *slelip* mutants may be affected in stomatal conductance. Gas exchange recordings of GH-grown plants were conducted at different light conditions during the day (Fig. 4). No major differences were observed in the stomatal conductance of *slelip* mutants vs M82 in the GH, where plants were subjected to ∼100 µmol photons m^-2^ s^-1^ (early morning) or 400-500 µmol photons m^-2^ s^-1^ (noon and afternoon) (Fig. 4a). Only a single difference, of higher conductance, was noted for the IK6 line around mid-day. Recordings were also conducted on a sunny day, while plants were continuously exposed to SL for 6 h. Under these conditions, stomatal conductance was higher in the morning, but no differences between *slelip* mutants and M82 were observed (Fig. 4b). Under both GH and SL conditions, no differences were apparent in the CO_2_ assimilation rates of M82 and mutant plants along the day (Fig. 4c,d), aside from one higher value for IK6 in the GH during afternoon (Fig. 4c).

**Fig. 4.**
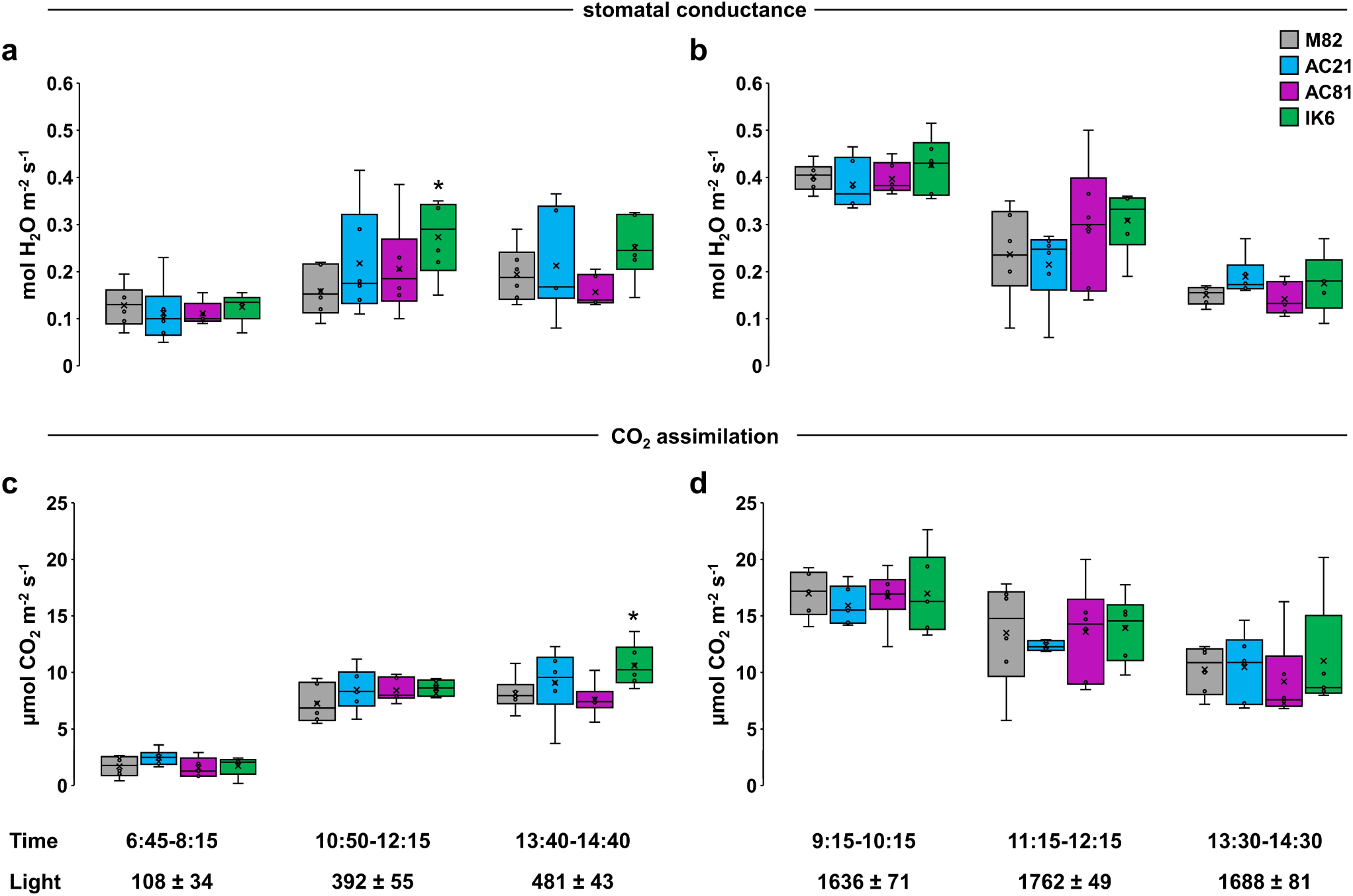
Stomatal conductance and CO_2_ assimilation are largely unaffected in *slelip* mutants at different times and light conditions. **(a, b)** Stomatal conductance and **(c, d)** CO_2_ assimilation of M82 and *slelip* mutants AC21, AC81, IK6. Gas exchange measurements were carried out on 3-months-old plants grown in a greenhouse under the greenhouse conditions (a, c), and under direct sunlight (b, d). The times of measurement and the light intensity ranges (µmol photons m^-2^ s^-1^) during the recordings are noted below the graphs. To depict the differences between conditions, the *y*-axis is shown in the same range. For each time point, n = 5-6 leaves per genotype, each from a different plant. Significant differences vs M82 are marked \**p* < 0.05 (Student’s T-test).

Finally, we tested whether *slelip* mutants may be affected during the initial response of stomata to light, by recording gas exchange during first exposure to light in the morning. Toward this, GH plants were brought into a dark room prior to dawn, and gas exchange recordings were conducted along with initial illumination (∼120 µmol photons m^-2^ s^-1^, white light) of the leaves. Remarkably, stomatal conductance in the three *slelip* mutant lines was reduced compared to M82 during most of the initial 25 min following the onset of illumination (Fig. 5).

**Fig. 5.**
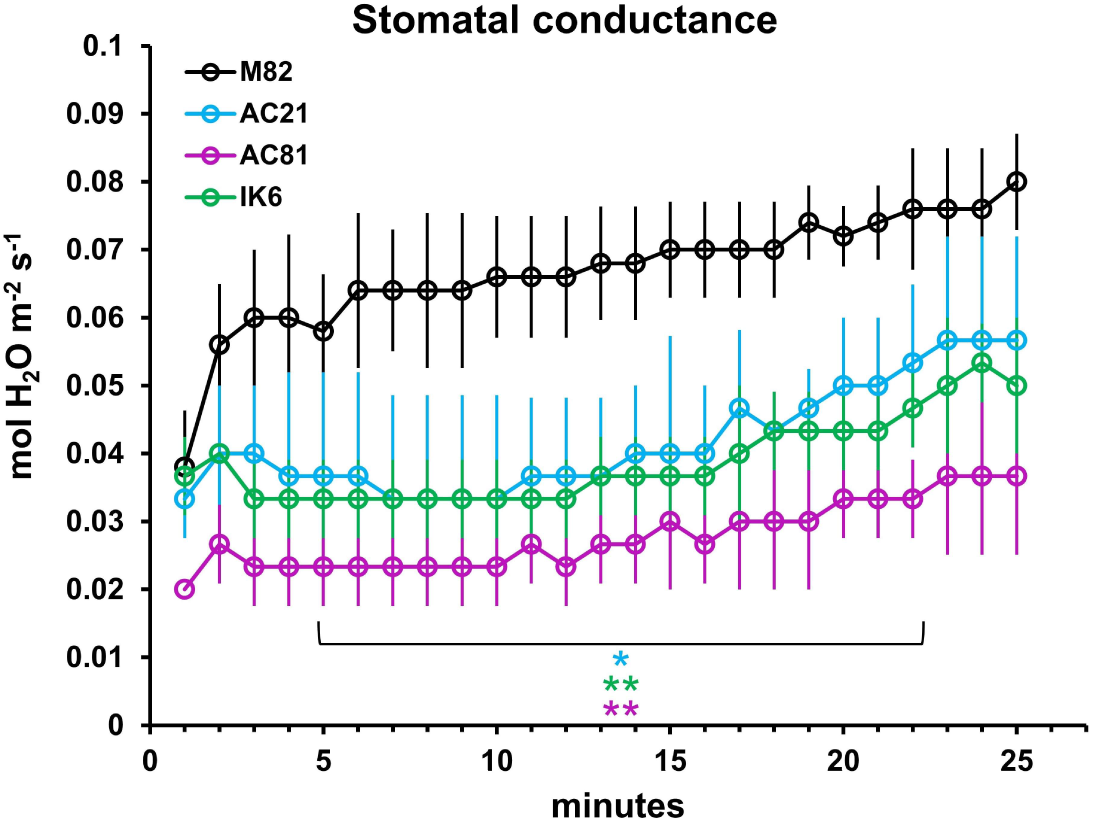
*slelip* mutants exhibit reduced stomatal conductance upon the transition to light. Greenhouse-grown plants were placed in a dark room prior to dawn. Each plant was exposed to white LED light (∼120 µmol photons m^-2^ s^-1^) while simultaneously recording stomatal conductance every minute for 25 min. All measurements were recorded in the timeframe between 6:00-8:00. Data are means ± SD of n = 4 plants (M82) and 3 plants for each mutant (AC21, AC81, IK6). The values of all mutant lines were significantly lower vs M82 between minutes 5 to 22, \**p* < 0.05, \*\**p* < 0.01 (Student’s T-test).

## Discussion

Plant ELIPs are most famed for a role in photoprotection. However, no clear detailed mechanistic information is available for ELIPs’ function in photoprotection, and this role is still under debate. Here, we studied ELIPs in tomato plants; on one hand characterizing *slelip* mutants, and on the other following the expression patterns of Venus or SlELIP1-Venus fused to the native *SlELIP1* promoter. Our main findings: **(1)** Investigation of *slelip* mutants revealed no indications for heightened sensitivity to photoinhibition, and do not conform with a general role of photoprotection. **(2)** *pSlELIP1* confers preferential expression in the leaf epidermis, most prominently in stomatal guard cells (GC), and the native SlELIP1 protein was confirmed to be considerably enriched in the epidermis. **(3)** All three mutants, two *slelip1* and one double mutant, exhibited reduced stomatal conductance upon the onset of illumination. Based on these, we propose that SlELIP1 has a role in the response of stomatal GC to light.

The *SlELIP1* promoter has been studied earlier and described as a fruit-specific promoter [at the time named ‘SGN-U144410’, (Estornell *et al*., 2009)]. The promoter fused to the GUS reporter showed that it confers increasing expression during fruit development with a maximal level in red fruit, yet very low expression in non-fruit tissues including leaves (Estornell *et al*., 2009). *pSlELIP1* used here was based on the *ELIP* ‘full-length’ promoter sequence (*S. lycopersicum* cv. Yalf) described by (Timerbaev & Dolgov, 2019). Detailed truncation analysis using promoter-GUS fusions revealed various positive and negative regulatory motifs, light-responsive elements, as well as ones involved in circadian and hormone response. Under the full-length *ELIP* promoter, relatively weak GUS expression was observed in leaves and immature fruit, while strong expression was visible in fruits at other stages (Timerbaev & Dolgov, 2019). The weak (GUS) expression in leaves can be explained by our confocal microscopy results showing preferential expression in the leaf epidermis, which represents a relatively thin fraction of the leaf, and would likely not be prominently detected by GUS staining of whole leaves.

In addition to our findings in tomato, data from other plants point to the expression of ELIPs in the epidermis and also in GC. Interestingly, a search for epidermal-selective promoters in *Medicago truncatula*, conducted by comparative transcriptomic analysis, discovered the *MtELIP* promoter as one with strong preferential expression in the upper epidermis (Cui *et al*., 2022). Notably, an earlier transcriptome study of GC in *A. thaliana* showed that the expression of *AtELIP1* is ∼32-fold higher in GC vs the leaf (Bates *et al*., 2012). Evidence for ELIPs’ preferential expression in the leaf epidermis in three species representing the Solanaceae, Fabaceae, and Brassicaceae families, suggests that this is a general trait of angiosperms (at least dicotyledons). Studies of *A. thaliana* have noted that on the protein level ELIPs do not accumulate in mature leaves (Casazza *et al*., 2005), or that they are lacking altogether under lab/indoor as opposed to field conditions (Mishra *et al*., 2012; Flannery *et al*., 2021). Considering that ELIPs are preferentially found in the epidermis, and that this tissue is a relatively small fraction of the leaf, the presence of ELIPs in the epidermis may be missed when analyzing extracts from whole leaf tissues. Likewise, ELIPs’ preferential expression in the leaf epidermis may explain why its putative roles in mesophyll photosynthesis and specifically in photoprotection have remained ambiguous. We note that perhaps ELIPs do have a role related to photoprotection in epidermal GC or other tissues, or at different plant developmental stages, yet this will require further detailed investigation.

The finding that SlELIP1 affects the initial response of stomata to light is intriguing. Numerous pathways operate in stomatal GC, finely regulating their function at different times and conditions. These include responses to light, CO_2_, hormones [abscisic acid (ABA), jasmonic acid], metabolites and more (Assmann & Jegla, 2016; Zhang *et al*., 2018; Lawson & Vialet-Chabrand, 2019; Matthews *et al*., 2020; Yang *et al*., 2020). The finding that *slelip* mutants exhibited reduced stomatal conductance at the onset of illumination in the morning, suggests that SlELIP1 positively affects stomatal opening upon the transition to light. A notable recent finding is that expression of the light-signaling ELONGATED HYPOCOTYL5 (HY5) transcription factor specifically in GC of *A. thaliana* resulted in increased Gs and transpiration (Kelly *et al*., 2023). HY5 is a positive regulator of *ELIP* genes, shown under different conditions in *A. thaliana* (Harari-Steinberg *et al*., 2001; Kleine *et al*., 2007; Hayami *et al*., 2015; Cañibano *et al*., 2021). Considering ELIPs expression and proposed function in GC, it will be interesting to investigate whether the effects of GC-expressed HY5 on stomatal function are related to ELIPs.

Light-induced stomatal opening occurs via two main described pathways, in response to blue or red light. The blue light response is a GC-specific pathway that promotes stomatal opening in the morning and saturates at low intensities (Kinoshita *et al*., 2001; Horrer *et al*., 2016; Inoue *et al*., 2020; Rovira *et al*., 2024). Alternatively, the response of GC to red light has been generally attributed to mesophyll photosynthesis, accordingly saturating at higher light intensities (Matthews *et al*., 2020; Flütsch & Santelia, 2021). Yet, there is also evidence that points to direct sensing of red light by GC (Zhu *et al*., 2020; Li *et al*., 2023; Hayashi *et al*., 2024). GC chloroplasts are essential for stomatal opening (Suetsugu *et al*., 2014), yet the involved mechanisms are not fully understood. ELIP is a chlorophyll-binding protein, shown here to be considerably enriched in the epidermis (as opposed to other LHC and LHC-like proteins), and specifically in GC, and to affect morning stomatal conductance. ELIP may therefore provide a new connection between light, GC chloroplasts and stomatal function. Deciphering the precise role(s) of ELIPs in GC still requires further detailed study. Yet, this is the first time that ELIPs are implicated in stomatal function, opening a new and exciting research direction for ELIPs’ long-sought roles.

## Acknowledgements

We thank Tal Dahan-Meir, Shiri Barad-Kotler and Ziva Amsellem (Weizmann Institute) for their help in generating the CRISPR/Cas9 mutants, and Smadar Greenstein (Volcani Institute) for her assistance with experiments. We thank Zach Adam (Hebrew University of Jerusalem), Ziv Reich, Reinat Nevo, and Yishai Levin (Weizmann Institute), and Yuval Cohen, Hagai Yasuor and Yoram Eyal (Volcani Institute) for their valuable advice and discussions. This work was supported by the Israel Science Foundation grant no. 1377/18 (to DC).

## Competing interests

None declared.

## Author contributions

DC, DB, MC, IK and VT designed the research. All authors performed experiments and collected data. DB, MC, VT, IK, PR and DC analyzed the data. DB and MC contributed equally. DC wrote the manuscript. All authors reviewed and approved the final version.

## Data availability

The mass spectrometry proteomics data have been deposited to the ProteomeXchange Consortium via the PRIDE (Perez-Riverol *et al*., 2025) partner repository with the dataset identifier PXD061612.

## Notes

### Competing Interest Statement

The authors have declared no competing interest.

