## Supporting Information for "ELIP is preferentially expressed in stomatal guard cells and has a role in stomatal opening upon transition to light"

**Table S1. Sequences of primers and gRNAs.**

| **Gene name and number** | | **Protein name** | **Primer/gRNA sequence** |
|---|---|---|---|
| ***Gene amplification and sequencing*** | | | |
| *SlELIP1* | Solyc09g082690 | early light-induced protein 1 | F: GATGCGCATTTACACTATTATC<br>R: GAAGAAACTATGCTATCTGATTACC |
| *SlELIP2* | Solyc09g082700 | early light-induced protein 2 | F: GTTCGTGTTAAGTGTATGGCTGAG<br>R: CCATTACTCATTAAGTGAAGGCA |
| ***qRT-PCR*** | | | |
| *SlELIP1* | Solyc09g082690 | early light-induced protein 1 | F: GTTCCTAGCTGTTACTTGCCAC<br>R: GGTGCTTGGTTTTGGCTTAGC |
| *SlELIP2* | Solyc09g082700 | early light-induced protein 2 | F: ATGTTAAAGGAGCTGGGCTT<br>R: CCATTACTCATTAAGTGAAGGCA |
| *SlCAC* | Solyc08g006960 | reference gene | F: CCTCCGTTGTGATGTAACTGG<br>R: ATTGGTGGAAAGTAACATCATCG |
| *SlSAND* | Solyc03g115810 | reference gene | F: TTGCTTGGAGGAACAGACG<br>R: GCAAACAGAACCCCTGAATC |
| ***gRNAs used for CRISPR/Cas targeting*** | | | |
| *SlELIP1* | Solyc09g082690 | gRNA ‘A’ | attgTAGCAGAGTAATCAGTTGA |
| *SlELIP1* | Solyc09g082690 | gRNA ‘C’ | attgACCTTAGCAGGTGTTGGCTT |
| *SlELIP1* & *SlELIP2* | Solyc08g006960<br>Solyc09g082700 | gRNA ‘I’ | attgATTTGTAGCAGCCATTGGTA |
| *SlELIP1* & *SlELIP2* | Solyc08g006960<br>Solyc09g082700 | gRNA ‘K’ | attgAGGCGTGGCAAGTAACAGCT |

**Liquid chromatography mass spectrometry methods**

*Sample preparation.* Fruit or leaf powder was suspended in lysis buffer containing 5% SDS in 50 mM Tris-HCl pH 7.4. Lysates were incubated at 96°C for 5 min, followed by six cycles of 30 s of sonication (Bioruptor Pico, Diagenode, USA). Protein concentration was measured using the BCA assay (Thermo Scientific, USA). 20 µg of total protein was reduced with 5 mM dithiothreitol and alkylated with 10 mM iodoacetamide in the dark. Each sample was loaded onto an S-Trap microcolumn (Protifi, USA) according to the manufacturer's instructions. After loading, samples were washed with 90:10% methanol/50 mM ammonium bicarbonate. Samples were then digested with trypsin (1:50 trypsin:protein ratio) for 1.5 h at 47°C. The digested peptides were eluted using 50 mM ammonium bicarbonate. Trypsin (1:50 trypsin:protein ratio) was added to this fraction and incubated overnight at 37°C. Two more elutions were made using 0.2% formic acid and 0.2% formic acid in 50% acetonitrile. The three elutions were pooled together and vacuum-centrifuged to dryness. Samples were kept at -80°C until further analysis.

*Liquid chromatography.* ULC/MS grade solvents were used for all chromatographic steps. *Fruit samples:* Each sample was loaded using split-less nano-Ultra Performance Liquid Chromatography (10 kpsi nanoAcquity; Waters, Milford, MA, USA). The mobile phase was: A) $H_2O$ + 0.1% formic acid and B) acetonitrile + 0.1% formic acid. Desalting of the samples was performed online using a Symmetry C18 reversed-phase trapping column (180 µm internal diameter, 20 mm length, 5 µm particle size; Waters). The peptides were then separated using a T3 HSS nano-column (75 µm internal diameter, 250 mm length, 1.8 µm particle size; Waters) at 0.35 µL/min. Peptides were eluted from the column into the mass spectrometer using the following gradient: 4% to 35%B in 105 min, 35% to 90%B in 5 min, maintained at 95% for 5 min and then back to initial conditions. *Leaf samples:* Dry digested samples were dissolved in 97:3% $H_2O$/acetonitrile + 0.1% formic acid. Each sample was loaded using nanoflow liquid chromatography (nanoElute2; Bruker Daltonics). The mobile phase was as above. The peptides were separated using an Aurora column (75µm ID x 25cm, IonOpticks) at 0.3 µL/min. Peptides were eluted from the column into the mass spectrometer using the following gradient: 2% to 35%B in 60 min, 35% to 95%B in 0.5 min, maintained at 95% for 6.77 min and then back to initial conditions.

*Mass Spectrometry. Fruit samples:* The nanoUPLC was coupled online through a nanoESI emitter (10 µm tip; New Objective; Woburn, MA, USA) to a quadrupole orbitrap mass spectrometer (Q Exactive HFX, Thermo Scientific, for discovery analysis, and Q Exactive Plus for targeted analysis) using a FlexIon nanospray apparatus (Proxeon). For discovery analysis data were acquired in data dependent acquisition (DDA) mode, using a Top10 method. MS1 resolution was set to 120,000 (at 200 m/z), mass range of 375-1650 m/z, AGC target of 3e6 and maximum injection time was set to 60 msec. MS2 resolution was set to 15,000, quadrupole isolation 1.7m/z, AGC of 1e5, dynamic exclusion of 30 sec and maximum injection time of 60 msec. For targeted analysis, data were acquired in parallel reaction monitoring (PRM) mode. MS1 resolution was set to 120,000 (at 200 m/z), mass range of 375-1650 m/z, AGC target of 3e6 and maximum injection time was set to 60 msec. MS2 resolution was set to 15,000, quadrupole isolation 1.7 m/z, AGC of 2e5, and maximum injection time of 100 msec. Inclusion list of peptides to be monitored included 15 peptides that are either unique or common to SlElip1 and SlElip2. *Leaf samples:* The nanoUPLC was coupled online to a timsTOF PRO mass spectrometer (Bruker Daltonics). Data was acquired in parallel accumulation–serial fragmentation combined with data independent acquisition (DIA-PASEF) mode (Meier *et al.*, 2018). For ion mobility 1/K0 range was set to 0.64-1.48 Vs/cm$^2$, ramp time of 100 msec and estimated cycle time of 1.48 sec. 40 windows of 25 Da with 1 Da overlap were set over a range of 262.4-1223.4 Da.

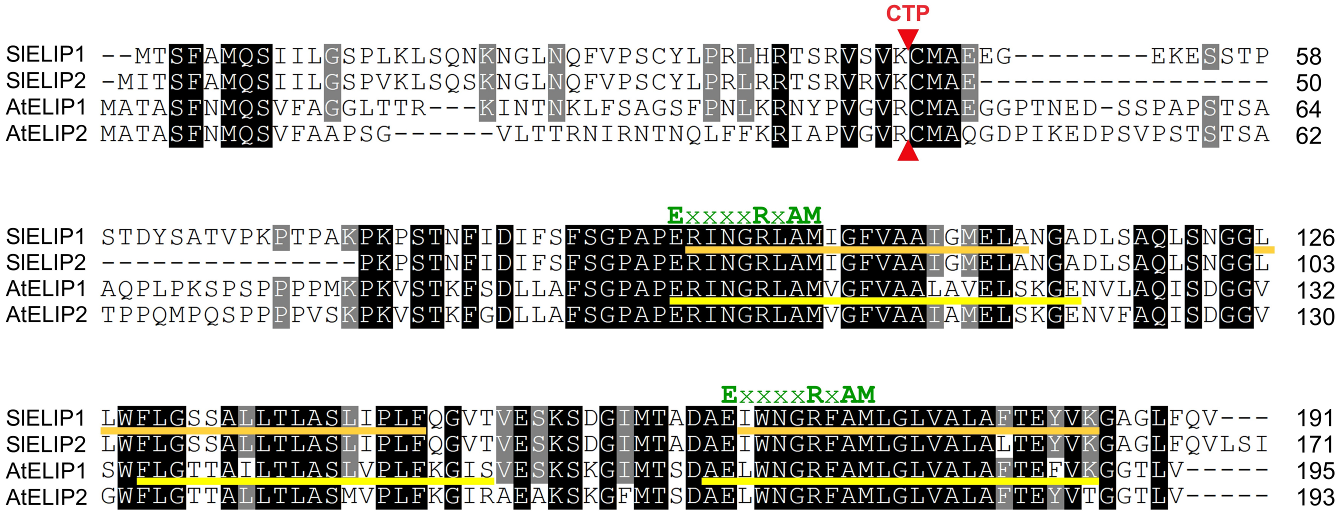


**Fig. S1. Protein sequence alignment of tomato and Arabidopsis ELIPs.** Alignment of SlELIP1 (Solyc09g082690) and SlELIP2 (Solyc09g082700) polypeptides with AtELIP1 (At3g22840) and AtELIP2 (At4g14690). Red triangles mark the predicted cleavage site of the chloroplast transit peptide (CTP). Orange lines indicate the predicted transmembrane regions in SlELIPs, and yellow lines the transmembrane spans of AtELIPs as shown in (Heddad & Adamska, 2000). The conserved chlorophyll-binding LHC motifs, ‘ExxxxRxAM’, in the first and third helices are noted above the sequence in green.

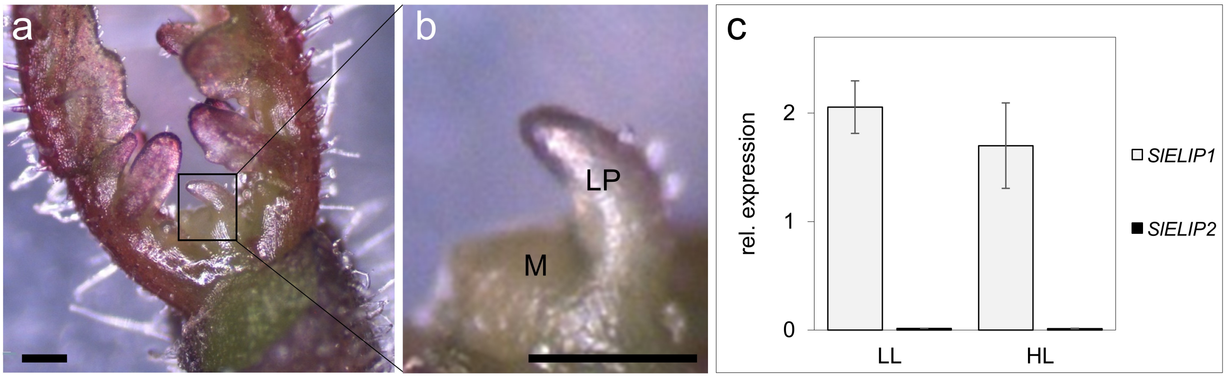


**Fig. S2. Expression of *SlELIP* genes in the shoot apex. (a)** The shoot apex region (black box) of a 10-day old tomato seedling. **(b)** Zoom-in view showing the shoot apical meristem (M) and leaf primordia (LP). **(c)** Expression of the *SlELIP1* and *SlELIP2* genes relative to the tomato *CAC* gene in seedlings germinated and grown at low light (LL; ~250 µmol photons $m^{-2}$ $s^{-1}$) or high light (HL; ~750 µmol photons $m^{-2}$ $s^{-1}$). Bars denote SE between 3 technical repetitions.

Method: For each light condition, 20-30 seedlings grown at the abovementioned light intensities, were collected and immediately transferred to chilled acetone. Samples were vacuum infiltrated for 30 min, and acetone replaced after 15 min. Samples were then kept at -80°C prior to dissection. Before dissection, samples were transferred to -20°C for 1 h, and then dissected under the binoculars in chilled acetone using a surgery knife (K2-5000; Katena). Acetone was evaporated at room temperature and RNA was then extracted by the PicoPure isolation kit (Zymo Research). cDNA was prepared from the two RNA preparations using the M-MLV reverse transcriptase (ThermoFisher).

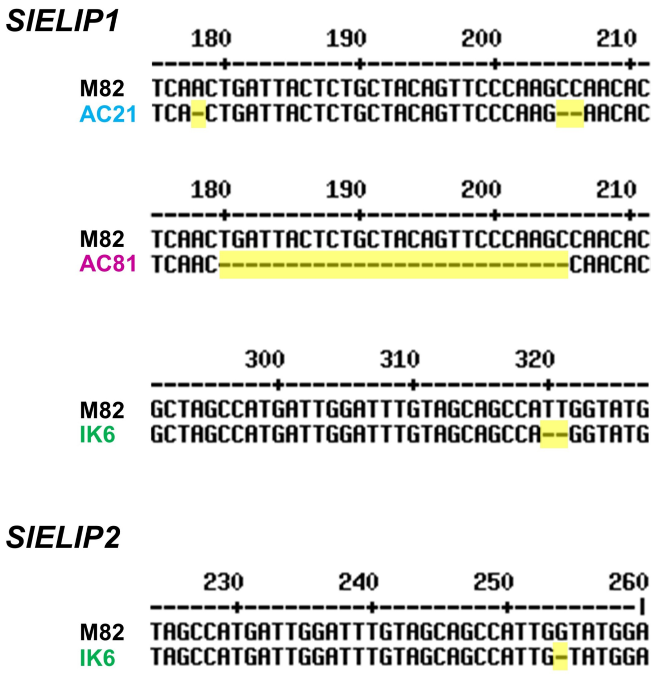


**Fig. S3. Sequence deletions in *SlELIP* genes.** 'AC21' and 'AC81' are lines mutated in *SlELIP1* and 'IK6' is a line mutated in both *SlELIP1* and *SlELIP2*. Nucleotide deletions (vs M82) are highlighted in yellow.

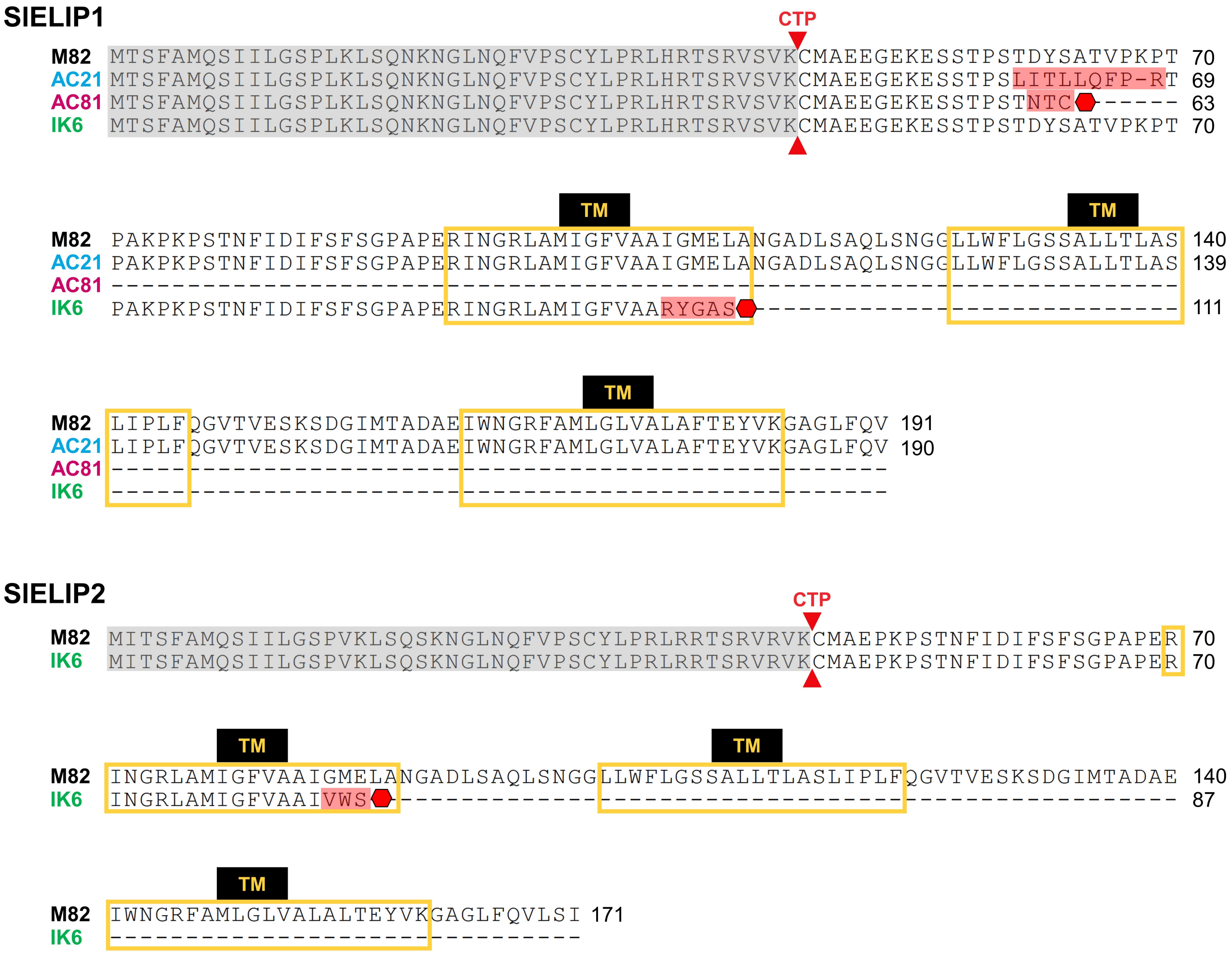


**Fig. S4. Sequence alignments of SlELIP polypeptides in M82 and in mutant lines (putative).** Red triangles mark the predicted chloroplast transit peptide (CTP) cleavage sites; transmembrane (TM) regions are shown by yellow boxes; altered sequences in mutants are highlighted in red; premature stop codons are depicted as red hexagons.

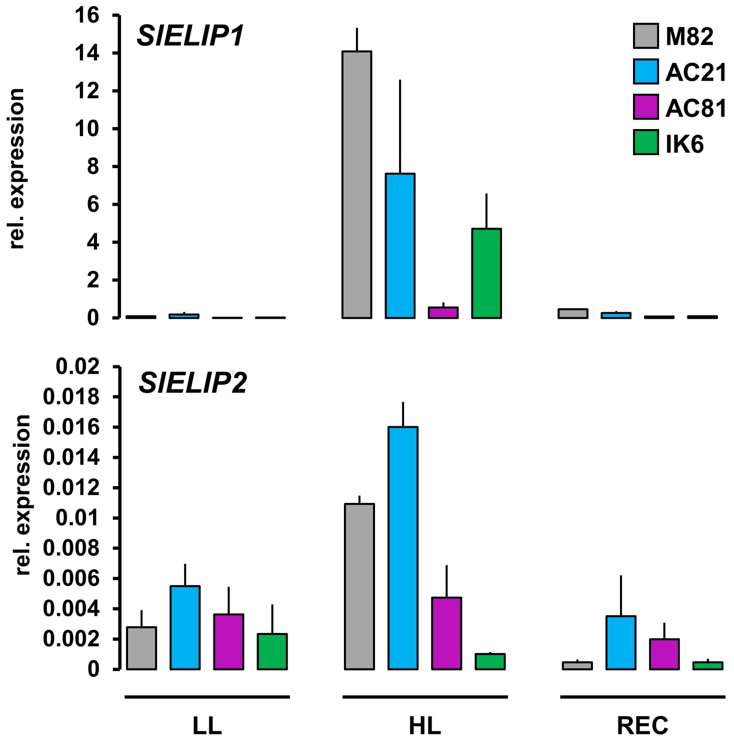


**Fig. S5. *SlELIP1* and *SlELIP2* expression in mutant lines vs M82.** Expression was probed in 6-week-old plant adapted to low light (LL), after exposure to 6 h of high light (HL), and following 24 h of recovery (REC) at initial LL conditions. Values shown are means ± SD of three biological (plant) replicates analyzed by qRT-PCR, normalized to two reference genes (*SlSAND* and *SlCAC*).

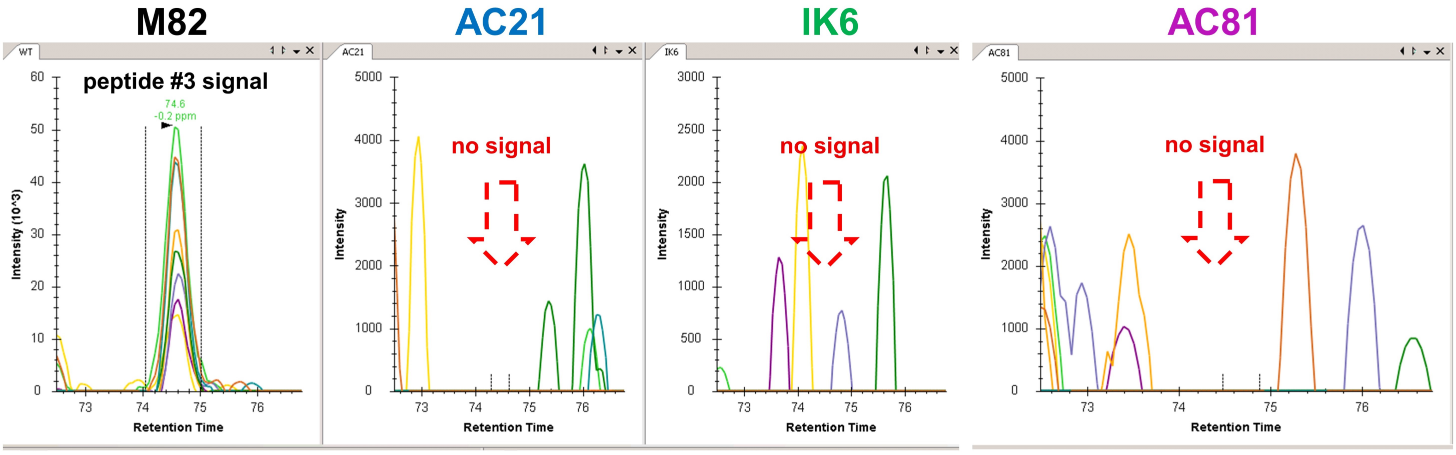


**Fig. S6. SlELIP peptide detection by targeted LC-MS/MS of orange fruit.** Lysates from orange fruit pericarp of M82, AC21, AC81 and IK6 were subjected to tryptic digestion and targeted LC-MS/MS analysis (Parallel Reaction Monitoring experiment), while monitoring SlELIP peptides. Extracted fragment ion chromatograms (generated by the Skyline algorithm) are shown for fragments of the SlELIP peptide 'SDGIMTADAEIWNGR' (peptide #3, Fig. 1b) in M82; each colored line corresponds to a different fragment. Dashed-line arrows in AC21, AC81 and IK6 indicate the expected retention time position for the eluted peptide, based on the chromatographic elution of the peptide in M82.

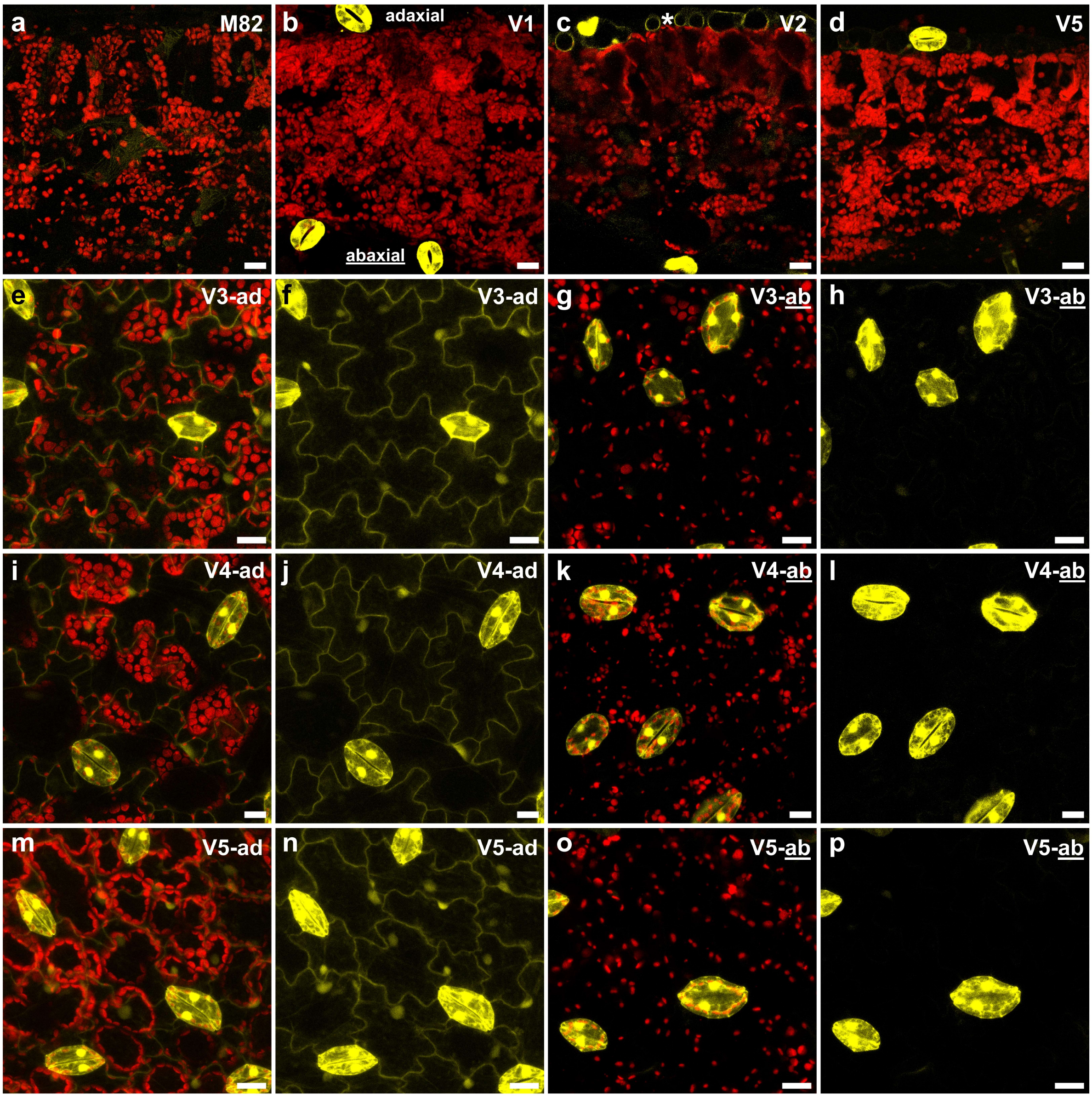


**Fig. S7. Venus fluorescence in cotyledons of *pSlELIP1*::Venus ('V') lines.** Confocal microscope images showing chlorophyll (Chl; red) and Venus-YFP (yellow) fluorescence in cross-sections and in adaxial ('ad') and abaxial ('ab') surfaces of cotyledons from 10-12-day-old seedlings germinated at low light (80-100 μmol photons $m^{-2}$ $s^{-1}$). **(a-d)** Cross-sections showing Chl and Venus in **(a)** M82, and in **(b-d)** V lines; asterisk in **(c)** denotes the visible Venus expression in adaxial pavement cells. Views of the adaxial surfaces of V lines show: **(e, i, m)** both Chl and Venus, or **(f, j, n)** Venus. Views of the abaxial surfaces of V lines show: **(g, k, o)** both Chl and Venus, or **(h, l, p)** Venus. The brightness of images showing only Venus **(f, h, j, l, n, p)** was increased to visualize the comparatively lower fluorescence of pavement cells (where visible).

Cross-section images are 9-10 µm thick (*Z*-thickness); surface images are 28-32 µm thick (encompassing the epidermal layer and part of the mesophyll in adaxial or abaxial surfaces). Scale bars = 20 µm.

**Table S2.** Ratio of light-harvesting (chlorophyll *a*/*b* binding) and light-harvesting-like proteins in the upper epidermis (E) vs leaf (L) from M82 plants exposed to high light[†]

| Gene ID | Protein accession | Protein name[1] | Ratio E/L[2] | #peptides | iBAQ-E[3] | iBAQ –L |
|---|---|---|---|---|---|---|
| Solyc09g082690 | Q6QDC5 | ELIP1 | 6.29 | 6 | 4.17E+03 | 6.86E+02 |
| Solyc04g053130 | A0A3Q7G5P7 | SEP2 | 2.11 | 3 | 1.54E+03 | 8.41E+02 |
| Solyc03g005770 | A0A3Q7G195 | CAB | 1.90 | 7 | 8.83E+04 | 6.26E+04 |
| Solyc04g010190 | A0A3Q7G0G3 | CAB | 1.70 | 2 | 3.51E+02 | 2.56E+02 |
| Solyc05g056070 | A0A3Q7GQW4 | CAB | 1.67 | 8 | 3.32E+04 | 1.86E+04 |
| Solyc03g005760 | A0A3Q7FCW3 | CAB | 1.66 | 1 | 5.49E+02 | 3.31E+02 |
| Solyc07g063600 | A0A3Q7HHU2 | CAB | 1.58 | 3 | 5.95E+04 | 4.30E+04 |
| Solyc10g007690 | P27522 | CAB8 | 1.56 | 10 | 6.87E+04 | 7.20E+04 |
| Solyc10g006230 | P10708 | CAB7 | 1.55 | 2 | 9.97E+02 | 6.20E+02 |
| Solyc01g105030 | P27524 | CAP10A (CP24) | 1.54 | 4 | 5.00E+04 | 4.54E+04 |
| Solyc12g006140 | A0A3Q7J1V3 | CAB | 1.54 | 6 | 9.47E+04 | 6.40E+04 |
| Solyc09g014520 | A0A3Q7I0X4 | CAB | 1.49 | 15 | 2.20E+05 | 1.56E+05 |
| Solyc03g115900 | A0A3Q7FU35 | CAB | 1.47 | 3 | 3.53E+04 | 2.43E+04 |
| Solyc06g063370 | Q00321 | CAB9 (CP29) | 1.42 | 13 | 2.23E+05 | 2.15E+05 |
| Solyc04g082930 | A0A3Q7GBC5 | CAB | 1.40 | 2 | 2.18E+02 | 1.40E+02 |
| Solyc04g071930 | A0A3Q7G7T5 | OHP2 | 1.38 | 5 | 2.08E+03 | 1.47E+03 |
| Solyc01g105050 | P27525 | CAP10B (CP24) | 1.35 | 4 | 6.33E+03 | 4.80E+03 |
| Solyc12g056620 | A0A3Q7JB63 | OHP2 | 1.33 | 2 | 3.80E+02 | 1.12E+02 |
| Solyc02g070940 | P07370 | CAB1B | 1.31 | 2 | 8.89E+04 | 6.77E+04 |
| Solyc06g060340 | P54773 | PSBS | 1.30 | 7 | 8.89E+04 | 6.97E+04 |
| Solyc12g009200 | A0A3Q7J3F7 | CAB | 1.30 | 4 | 1.79E+03 | 1.17E+03 |
| Solyc07g022900 | A0A3Q7HA75 | CAB | 1.19 | 4 | 1.58E+03 | 1.16E+03 |
| Solyc07g047850 | P14278 | CAB4 | 1.14 | 1 | 1.33E+02 | 1.17E+02 |
| Solyc12g011280 | A0A3Q7JSZ2 | CAB | 0.99 | 3 | 8.18E+02 | 9.07E+02 |
| Solyc01g107660 | A0A3Q7ESD9 | SEP1 | 0.80 | 1 | 1.99E+02 | 2.47E+02 |
| Solyc06g069730 | Q7M1K8 | CAB | 0.73 | 1 | 1.14E+02 | 1.55E+02 |
| Solyc08g007180 | A0A3Q7IEJ1 | LIL3 | 0.72 | 4 | 3.79E+02 | 5.51E+02 |
| Solyc08g077880 | A0A3Q7INJ8 | LIL3 | 0.62 | 10 | 2.26E+03 | 3.51E+03 |

[†]The full list of (~7500) proteins identified by discovery LC-MS/MS is provided in Table S3.

[1]Protein names: ELIP, early light-induced protein; SEP, stress-enhanced protein; CAB, chlorophyll *a/b* binding protein; OHP, one helix protein; LIL, light-harvesting-like.

[2]Ratio in epidermis vs leaf is listed in descending order.

[3]iBAQ, intensity-based absolute quantification.
